# Metabolism of FAD, FMN and riboflavin (vitamin B2) in the human parasitic blood fluke *Schistosoma mansoni*

**DOI:** 10.1101/2024.03.12.584659

**Authors:** Akram A. Da’dara, Catherine S. Nation, Patrick J. Skelly

## Abstract

Schistosomiasis is a parasitic disease caused by trematode worms of the genus *Schistosoma.* The intravascular worms acquire the nutrients necessary for their survival from host blood. Since all animals are auxotrophic for riboflavin (vitamin B2), schistosomes too must import it to survive. Riboflavin is an essential component of the coenzymes flavin mononucleotide (FMN) and flavin adenine dinucleotide (FAD); these support key functions of dozens of flavoenzymes. In this work we focus on the biochemistry of riboflavin and its metabolites in *Schistosoma mansoni*. We show that when schistosomes are incubated in murine plasma, levels of FAD decrease over time while the levels of FMN increase. We show that live schistosomes can cleave exogenous FAD to generate FMN and this ability is significantly blocked when expression of the surface ectoenzyme SmNPP5 is suppressed using RNAi. Recombinant SmNPP5 cleaves FAD with a Km of 178 ± 5.9 µM. The FAD-dependent enzyme IL-4I1 drives the oxidative deamination of phenylalanine to produce phenylpyruvate and H_2_O_2_ in the extracellular environment. Since schistosomes can be damaged by H_2_O_2_, we determined if SmNPP5 could impede H_2_O_2_ production by blocking IL-4I1 action *in vitro*. We found that this was not the case, suggesting that covalently bound FAD on IL-4I1 is inaccessible to SmNPP5. We also report here that live schistosomes can cleave exogenous FMN to generate riboflavin and this ability is significantly impeded when expression of a second surface ectoenzyme (alkaline phosphatase, SmAP) is suppressed. Recombinant SmAP cleaves FMN with a Km of 3.82 ± 0.58 mM. Thus, the sequential hydrolysis of FAD by tegumental ecto-enzymes SmNPP5 and SmAP can generate free vitamin B2 around the worms from where it can be conveniently imported by, we hypothesize, the recently described schistosome riboflavin transporter SmaRT. In this work we also identified *in silico* schistosome homologs of enzymes that are involved in intracellular vitamin B2 metabolism. These are riboflavin kinase (SmRFK) as well as FAD synthase (SmFADS); cDNAs encoding these two enzymes were cloned and sequenced. SmRFK is predicted to convert riboflavin to FMN while SmFADS could further act on FMN to regenerate FAD in order to facilitate robust vitamin B2-dependent metabolism in schistosomes.

## Introduction

Schistosomiasis is a parasitic disease caused by trematode worms (blood flukes) belonging to the genus *Schistosoma.* Over 200 million people are estimated to be infected with these parasites around the world; transmission has been reported from 78 countries and over 800 million live at risk of infection ^1,2^. Schistosomiasis can result in abdominal pain, diarrhea, and blood in the stool or urine. The disease is associated with disabling anemia, growth stunting and poor performance at school and at work ^3^. People can become infected when larval forms of the parasite (cercariae) that are released by freshwater snails penetrate the skin. Once inside the host’s body, cercariae transform into juvenile forms called schistosomula. These migrate through the bloodstream to the portal vein, where they mature into adults. Adult male and female worms mate in the blood vessels and the females release eggs. Some of the eggs leave the body in feces or in urine to continue the parasite’s lifecycle. Other eggs can become trapped in body tissues, provoking immune reactions that can lead to progressive damage to organs.

Schistosomes can live for many years within the vasculature of their hosts where they acquire all nutrients necessary for their survival and growth ^4–6^. In this work we focus on one key schistosome species infecting humans, *Schistosoma mansoni*; we examine extracellular and intracellular enzymes of these worms that are involved in the metabolism of riboflavin (vitamin B2) and its key metabolites, flavin mononucleotide (FMN) and flavin adenine dinucleotide (FAD).

Riboflavin is a water-soluble vitamin that is essential for the formation of the two key coenzymes noted above, FMN and FAD ^7^. These coenzymes support the function of dozens of flavoenzymes in a wide variety of organisms; they are critical for the metabolism of carbohydrates, lipids, and proteins and they function as electron carriers in many oxidation-reduction (redox) reactions such as those that drive energy production ^8^. One example of a flavoenzyme is interleukin 4 induced 1 (IL-4I1). This protein is secreted from myeloid cells as well as B and T lymphocytes. It performs oxidative deamination of phenylalanine to produce phenylpyruvate and H_2_O_2_ and has been shown to be involved in the fine control of B- and T-cell adaptive immune responses ^9,10^. Flavoproteins are also involved in the metabolism of other B vitamins such as B3 (niacin), B6 (pyridoxal), B9 (folate), and B12 (cobalamin).

Biosynthesis of riboflavin can take place in bacteria, fungi and plants, but not in animals ^11^. This means that schistosomes and their hosts must both obtain sufficient amounts of riboflavin in their diets to support the functions of their myriad flavoenzymes and thus maintain optimal health. Clinical symptoms of riboflavin deficiency in humans include inflammation, anemia and impaired nerve function ^12^.

Riboflavin transporter proteins in animal cell membranes act as conduits for riboflavin uptake ^13,14^. Using the sequence of the human riboflavin transporter RFVT2 as a guide, we queried schistosome genome databases to identify a riboflavin transporter homolog in *S. mansoni* designated SmaRT ^15^. This 531 amino-acid protein is predicted to possess 11 transmembrane domains and has been immunolocalized to the tegument and the internal tissues of the adult worms. Functional expression of SmaRT in CHO-S cells showed it to be a *bone fide* riboflavin transporting protein that functions in a sodium independent manner and over a wide range of pH values ^15^.

FMN and FAD do not readily cross cell membranes; extracellular FMN and FAD need to be cleaved to generate free riboflavin which can then be imported by cells ^16^. In work with human cells, it has been shown that FAD is hydrolyzed by the ecto-5’ nucleotidase CD73 to generate FMN ^16^. Then, FMN is hydrolyzed by an ecto-alkaline phosphatase to riboflavin, which can be efficiently imported into cells ^16^. We hypothesized that a similar sequential hydrolysis of extracellular FAD through FMN to riboflavin occurs in intravascular schistosomes. Such riboflavin could be imported into the worms by the riboflavin transporter SmaRT ^15^.

Analysis of the schistosome tegumentome has not identified a schistosome ecto-5’ nucleotidase (akin to CD73), so how the worms might cleave extracellular FAD to generate FMN, is uncertain ^17,18^. However, the worms do express a tegumental phosphodiesterase/pyrophosphatase designated SmNPP5 that we have recently shown can cleave the related metabolite NAD ^19,20^. SmNPP5-mediated cleavage of NAD generates NMN and AMP ^20^. SmNPP5 is a 458 amino acid, glycophosphatidylinositol (GPI)-linked glycoprotein that can additionally cleave ATP, ADP and ADPR ^21,22^. We hypothesize here that SmNPP5 can also hydrolyze FAD.

As noted earlier regarding work with human cells, extracellular FMN is hydrolyzed by an ecto-alkaline phosphatase to generate free riboflavin ^16^. We have cloned and characterized a *S. mansoni* tegumental ecto-alkaline phosphatase designated SmAP that could conceivably likewise cleave exogenous FMN to generate riboflavin in schistosomes ^23^. SmAP is a 536 amino acid, GPI-linked tegumental glycoprotein ^23,24^. We have expressed SmAP in CHO-S cells, purified the recombinant protein and shown that it can dephosphorylate several substrates including AMP, CMP, GMP and TMP ^25^. SmAP additionally hydrolyzes the bioactive lipid sphingosine-1-phosphate (S1P) ^25^ as well as the proinflammatory and prothrombotic polymer, polyphosphate (polyP) ^26^.

Both SmAP and SmNPP5 are highly expressed in the intravascular life stages of *S. mansoni,* and both have been identified in several tegumental proteomics studies ^17,27,28^. In addition, both proteins immunolocalize to the external parasite tegument where they could come into contact with, and act upon, host metabolites such as FAD and FMN ^23,24^.

Any riboflavin that is taken up by schistosomes via SmaRT would need to be converted to FMN and FAD and, in this work, we also perform *in silico* analysis to determine if schistosomes possess genes that could encode enzymes to facilitate these conversions and so promote robust central metabolism.

## Results

### Live schistosomes can cleave exogenous FAD and FMN

To examine global metabolic changes brought about by schistosomes on host plasma, adult worms were first incubated in murine plasma as described in Methods. Twenty and sixty minutes later, samples were collected and changes to the plasma metabolome were measured. **Figure 1** illustrates relative changes in three metabolites FAD (panel A), FMN (panel B) and riboflavin (panel C) from murine plasma which contained (+) or did not contain (-) adult schistosomes. **Figure 1A** shows that there is a significant drop in the level of FAD at both the 20- and the 60-minute time point in the plasma sample that contained worms, and this is accompanied by a significant increase in plasma FMN at both time points (shown in **Figure 1B**, P<0.05 in each case). There is no significant change in riboflavin levels detected between any of the samples, as shown in **Figure 1C**.

**Figure 1.**
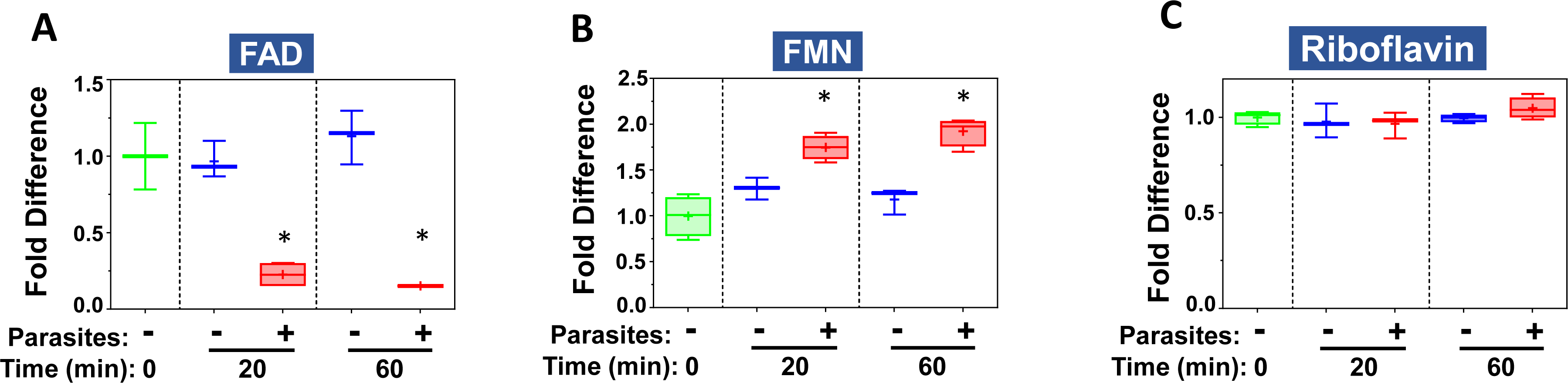
Box plots showing relative levels of FAD (**A**), FMN (**B**) and riboflavin (**C**) in murine plasma that either contained adult schistosomes (+, red) or did not contain schistosomes (-, blue) for the indicated time periods. * indicates statistically significant difference compared to the same time point lacking parasites; Welch’s two-sample t-test, P < 0.05. Each box bounds the upper and lower quartile, the line in each box is the median value and “+” signifies the mean value for the sample; error bars indicate the maximum (upper) and minimum (lower) distribution. Values obtained at zero time from plasma lacking worms (0, green symbols) are set at 1.

### Ectoenzyme SmNPP5 can hydrolyze FAD

We have cloned and expressed the schistosome ectoenzyme SmNPP5 in CHO-S cells ^25^. An aliquot of purified rSmNPP5 running at ∼55kDa and resolved by SDS-PAGE on a Coomassie-stained gel is shown in figure 2A (arrow), alongside molecular mass markers (right lane). Figure 2B shows that rSmNPP5 can cleave FAD to generate the fluorescent reaction product FMN; the reaction was carried out at two FAD concentrations 50µM (red line, figure 2B) and 500µM (blue line). Figure 2C shows that when the FAD cleavage product (FMN) is incubated with calf intestinal phosphatase (cip), free phosphate is released, and the amount released increases with increasing incubation time (from 0.5h to 5h). These data support the pathway presented in figure 2D in which rSmNPP5 cleaves FAD at the site depicted with a dashed pink line to generate FMN and AMP. The Michaelis Menten kinetics of SmNPP5-mediated cleavage of FAD was measured and, as shown in figure 2E, the Km of rSmNPP5 for FAD is 178 ± 5.9 µM.

**Figure 2.**
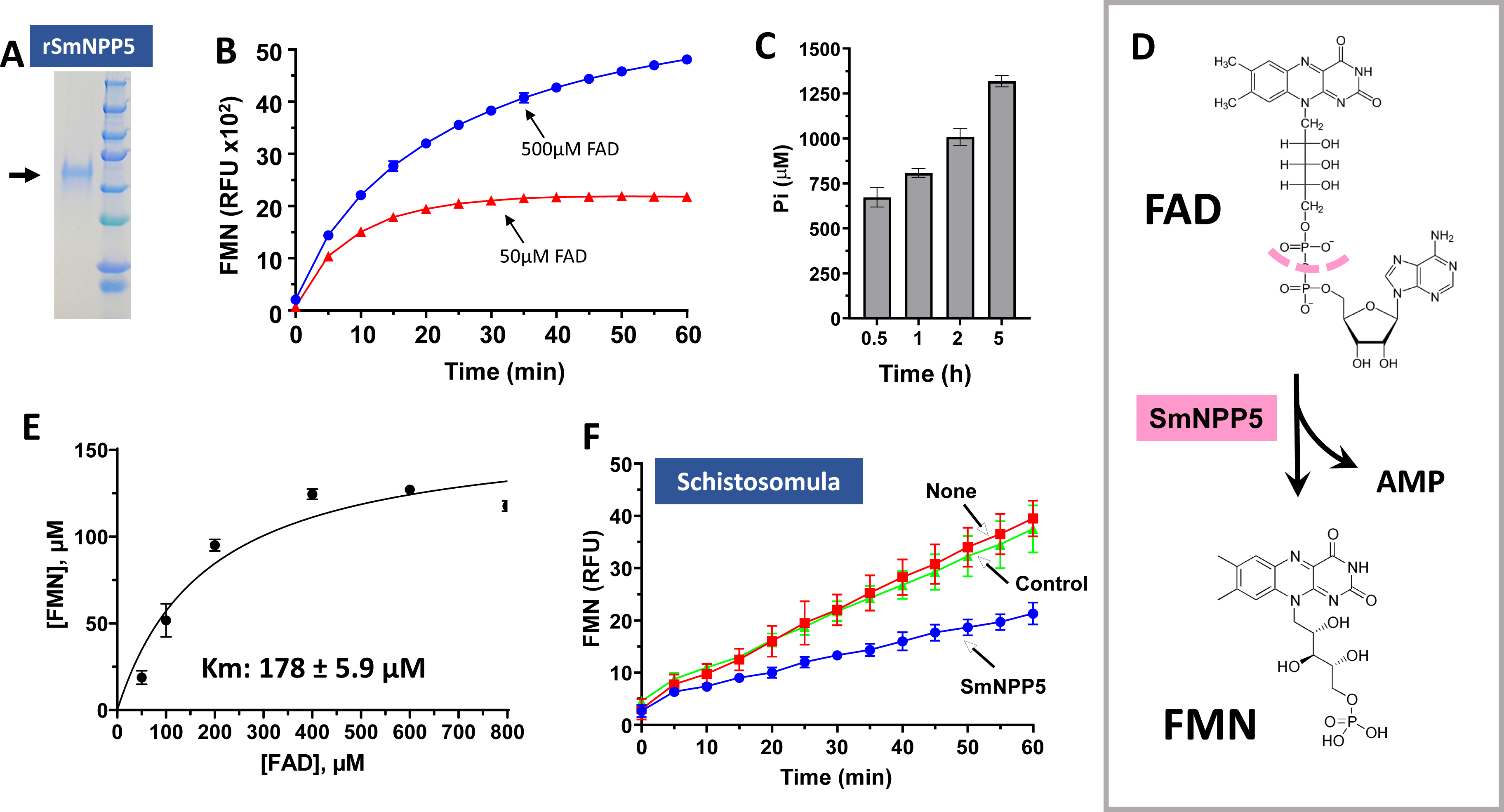
SmNPP5 can cleave FAD. **A**. An aliquot of purified rSmNPP5 resolved by SDS-PAGE is shown (arrow) alongside molecular mass markers (right lane). **B**. FMN generation (RFU, mean ± SD) by rSmNPP5 in assay buffer containing FAD as substrate (50 µM, red line or 500 µM, blue line), at the indicated time points. **C**. SmNPP5 was incubated with 500 µM FAD for 0.5, 1,2 or 5 h, as indicated, before an aliquot was withdrawn and calf intestinal phosphatase (cip) added. The figure shows phosphate generation (µM, mean ± SD) from each sample after 30 min. **D**. Depiction of SmNPP5-mediated cleavage of FAD (top) to generate FMN (bottom) and AMP. The pink dashed line indicates the site of cleavage. **E**. Michaelis-Menton plot of SmNPP5-mediated FAD cleavage kinetics; the Km of rSmNPP5 for FAD is 178 ± 5.9 µM, derived from three independent experiments. **F**. FMN generation (RFU, mean ± SD, n=5) by groups of ∼1,000 schistosomula seven days after treatment with siRNA targeting SmNPP5 (blue line) or an irrelevant siRNA (Control, green line) or no siRNA (None, red line) in the presence of FAD. Parasites treated with SmNPP5 siRNA generate significantly less FMN compared to either control (Two-way ANOVA, P < 0.01).

To test the hypothesis that tegumental SmNPP5 on live schistosomes is responsible for exogenous FAD cleavage, parasites were first treated with siRNAs targeting SmNPP5 or with control siRNAs or with no siRNA (None). Next, the ability of all worms to hydrolyze FAD was compared 7 days later. Figure 2F shows the results of this experiment and it is clear that worms whose SmNPP5 gene has been suppressed (blue line, figure 2F) cleave significantly less FAD compared to control worms treated either with the control siRNA (green line) or with no siRNA (None, red line; P<0.01 for the SmNPP5 suppressed group v either control).

### SmNPP5 does not inhibit the action of the FAD-dependent enzyme IL-4I1

The extracellular enzyme Interleukin 4 Induced 1 (IL-4I1) catalyzes the reaction depicted in figure 3A. As shown, this reaction generates hydrogen peroxide (H_2_O_2_, yellow highlight at right, figure 3A) which is potentially damaging to schistosomes ^29,30^. IL-4I1 activity requires FAD as a cofactor. Here we tested the ability of FAD-cleaving rSmNPP5 to inhibit the action of IL-4I1. Figure 3B shows that incubating rSmNPP5 with IL-4I1 does not block the enzyme’s activity (red line) compared to enzyme incubation with an irrelevant control protein (BSA, green line). Other controls (lower lines, figure 3B), as expected, elicit no activity, e.g., substrate incubated alone (Substrate) or rSmNPP5 incubated with substrate but without IL-4I1 (rSmNPP5). In separate experiments, we pre-incubated IL-4I1 with SmNPP5 for 1 h and then performed the enzyme assay; no inhibition was observed under these conditions too (data not shown).

**Figure 3.**
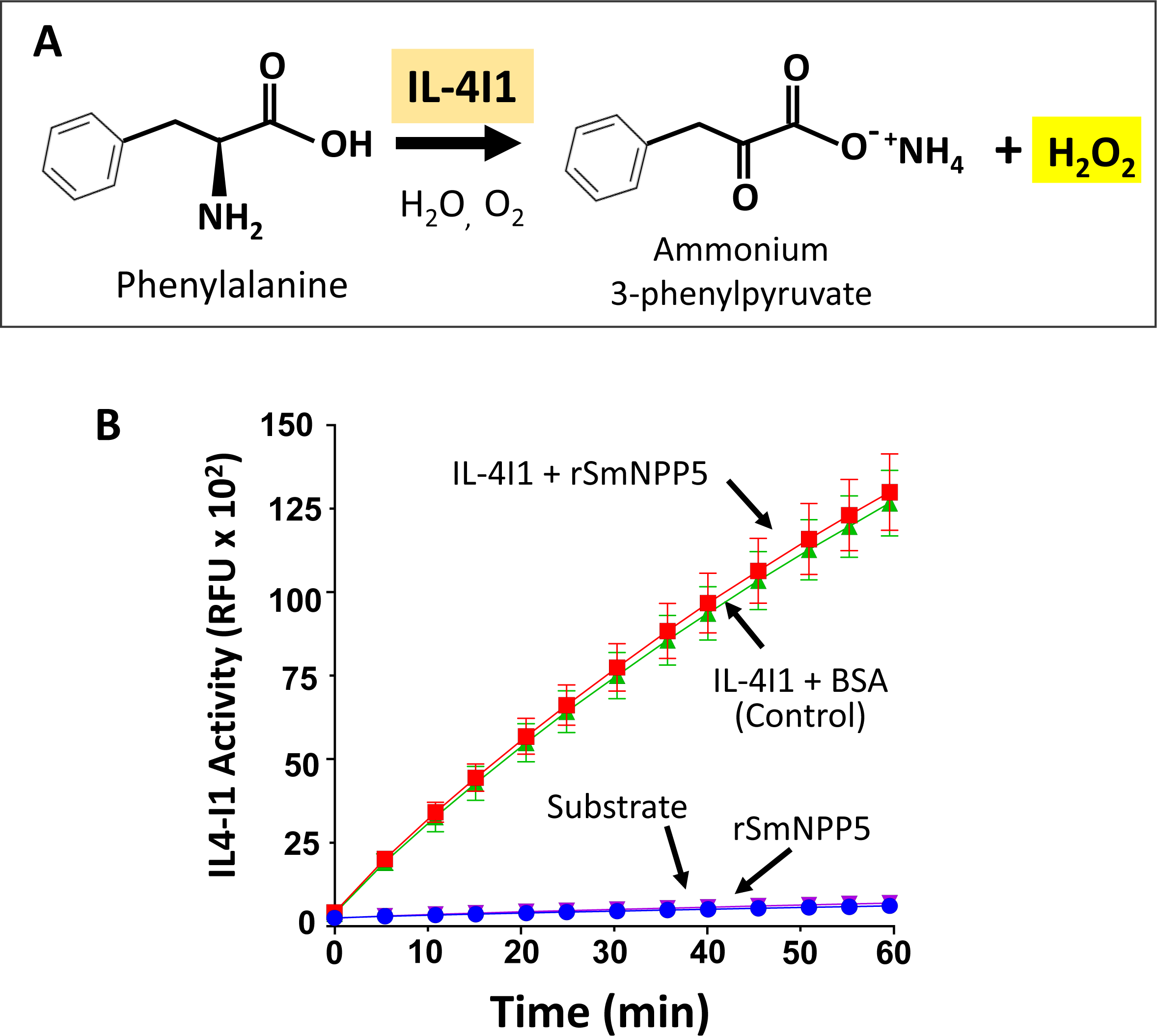
SmNPP5 does not inhibit IL-4I1 reactivity. **A**. The reaction driven by the FAD-dependent flavoenzyme IL-4I1, leading to the generation of H_2_O_2_ (yellow highlight, right), is depicted. **B**. IL-4I1 activity (RFU, mean ± SD, n=4) in the presence of either rSmNPP5 (red symbols) or control protein (BSA, green symbols). No significant difference in activity is detected under the two conditions. As expected, no activity is detected in the presence of substrate alone or rSmNPP5 alone (without IL-4I1), lower lines.

### Tegumental SmAP cleaves FMN

We have cloned and expressed the schistosome ectoenzyme SmAP in CHO-S cells ^25^. An aliquot of purified rSmAP running at ∼60 kDa and resolved by SDS-PAGE on a Coomassie-stained gel is shown in figure 4A (arrow), alongside molecular mass markers (right lane).To test the hypothesis that rSmAP can cleave FMN to generate riboflavin (RF) and phosphate (Pi), the purified enzyme was incubated with FMN, and any phosphate generated over time was measured. Figure 4B shows that Pi is indeed generated, with more being formed with prolonged incubation (from 0.5 to 2 h, black bars). No spontaneous breakdown of FMN is recorded over this time period (FMN alone, grey bars, Figure 4B). As shown in figure 4C, TLC confirms that incubating FMN (indicated by the blue arrowhead) with rSmAP (+ lane) yields riboflavin (RF, red arrowhead). Standards (FMN and RF) are resolved in the left TLC panel (figure 4C). These data support the pathway presented in figure 4D in which rSmAP cleaves FMN (blue arrowhead) at the site depicted with a dashed yellow line to generate RF (red arrowhead) and Pi. The Michaelis Menten kinetics of SmAP-mediated cleavage of FMN was measured and as shown in figure 4E, the Km of rSmAP for FMN is 3.82 ± 0.58 mM.

**Figure 4.**
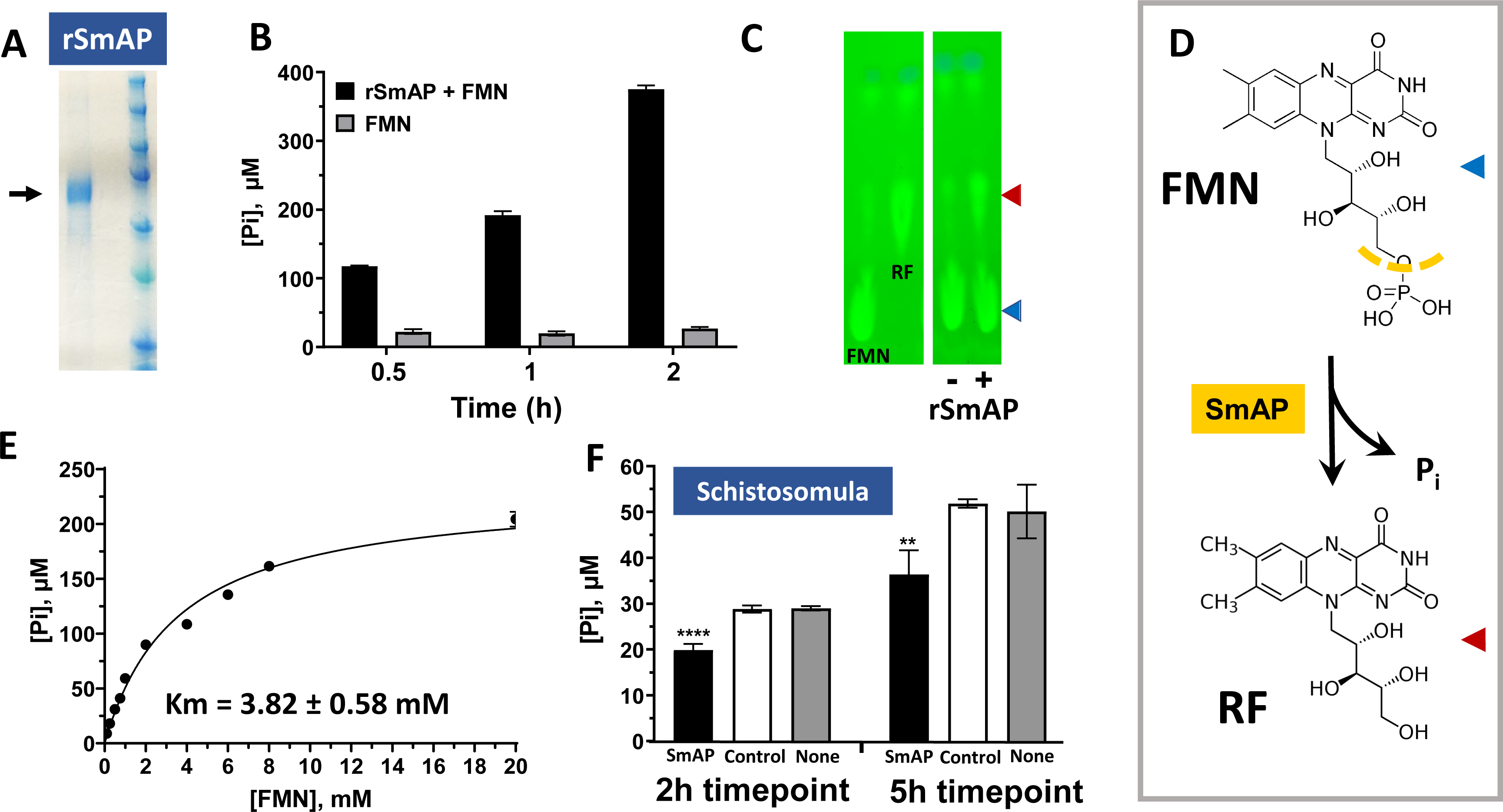
SmAP can cleave FMN. **A**. An aliquot of purified rSmAP resolved by SDS-PAGE is shown (arrow) alongside molecular mass markers (right lane). **B**. Phosphate (Pi) generation (µM, mean ± SD) by rSmAP in assay buffer containing FMN as substrate at the indicated time points (black bars). Essentially no Pi is generated when FMN is incubated without enzyme (grey bars). **C**. Thin layer chromatographic resolution of reaction products generated after incubating 1 mM FMN with (+) or without (−) rSmAP for 1h. The red arrowhead indicates reaction product riboflavin (RF). The blue arrowhead indicates the position of migration of FMN. Migration of standards is shown at left. **D**. Depiction of SmAP-mediated cleavage of FMN (top) to generate RF (bottom) and Pi. The yellow dashed line indicates the site of cleavage. The colored arrowheads indicate the molecules seen in the TLC image (depicted in panel C). **E**. Michaelis-Menton plot of SmAP-mediated FMN cleavage kinetics; the Km of rSmAP for FMN is 3.82 ± 0.58 mM, derived from three independent experiments. **F**. Pi generation (µM, mean ± SD, n=5) by groups of ∼1,000 schistosomula seven days after treatment with siRNA targeting SmAP (black bars) or an irrelevant siRNA (Control, white bars) or no siRNA (None, grey bars) in the presence of FMN. Parasites treated with SmAP siRNA generate significantly less Pi compared to either control (One-way ANOVA, **, p<0.01; ****p < 0.0001).

To test the hypothesis that tegumental SmAP on live schistosomes is responsible for exogenous FMN cleavage, parasites were first treated with siRNAs targeting SmAP or with control siRNAs or with no siRNA. Next the ability of all worms to hydrolyze FMN and to generate Pi was compared 7 days later. Figure 4F shows the results of this experiment and it is clear that worms whose SmAP gene has been suppressed (black bars, figure 4F) cleave significantly less NMN at both the 2 h and 5 h timepoints, compared to control worms treated either with control siRNA (white bars) or with no siRNA (None, grey bars; ** P<0.01, ****P<0.0001 v both controls).

### *In silico* analysis of vitamin B2 metabolizing cytosolic enzymes of *S. mansoni*

Our analysis suggests that any FAD in the vicinity of intravascular schistosomes could be cleaved by the tegumental ectoenzyme SmNPP5 to generate FMN. The liberated FMN could then be dephosphorylated by tegumental ectoenzyme SmAP to generate RF and this essential vitamin could then be conveniently taken up by the worms via the riboflavin transporter SmaRT to facilitate central metabolism. Since RF is key to the formation of two major enzyme coenzymes, FMN and FAD, any free RF that is taken in by schistosomes would need to be first phosphorylated internally to regenerate FMN and some of this FMN could be further metabolized to FAD. We examined schistosome DNA sequence databases to determine if the worms contain genes encoding cytosolic proteins that could drive the required reactions; RF to FMN and FMN to FAD.

In other systems a riboflavin kinase (RFK) converts RF to FMN. By blasting the human RFK protein sequence against schistosome sequence databases we identified a clear *S. mansoni* RFK homolog designated SmRFK. As detailed in Methods, the complete SmRFK coding sequence was amplified by PCR and the purified PCR product was cloned and sequenced. The full length schistosome SmRFK sequence was identical to a deposited sequence at the GenBank with the accession number XM_018794765. SmRFK potentially encodes a 154 amino acid protein with a predicted molecular mass of 17,681 Daltons and a pI of 7.91. Figure 5A shows an alignment of this SmRFK protein with counterparts from *S. haematobium* and *S. japonicum* as well as homologs from a variety of other organisms. Highly conserved residues found in all sequences are indicated with a star (*). Residues involved in flavin binding are colored red and indicated with (**$**), and those involved in ADP/ATP binding are colored blue and indicated with (**#**). The main residue (Thr^32^) involved in the binding of the magnesium ion (Mg^2+^) is colored green and indicated with (↓). Furthermore, SmRFK contains the highly conserved ^31^PTAN^34^ motif (boxed) that is found in all described RFK enzymes.

**Figure 5.**
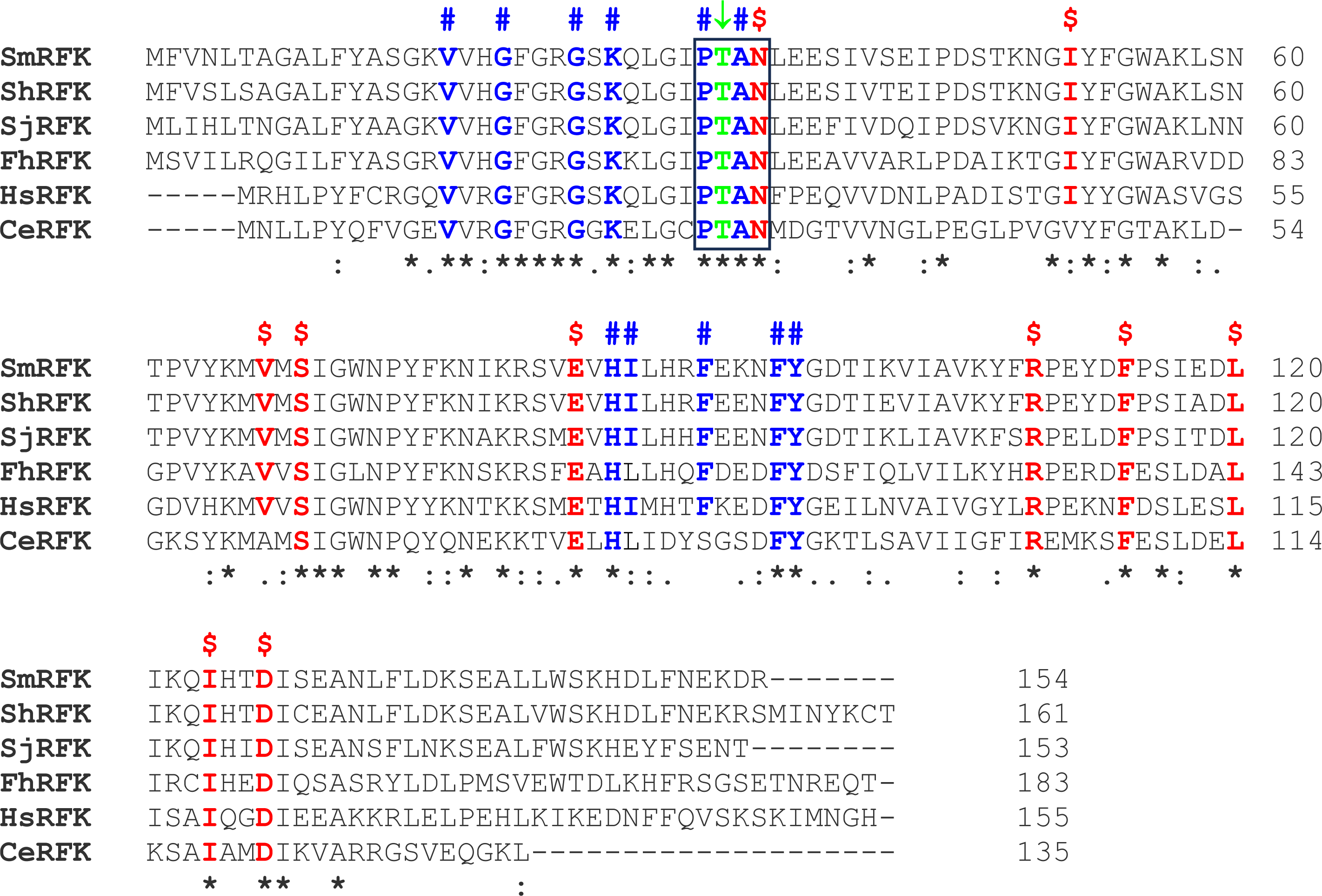
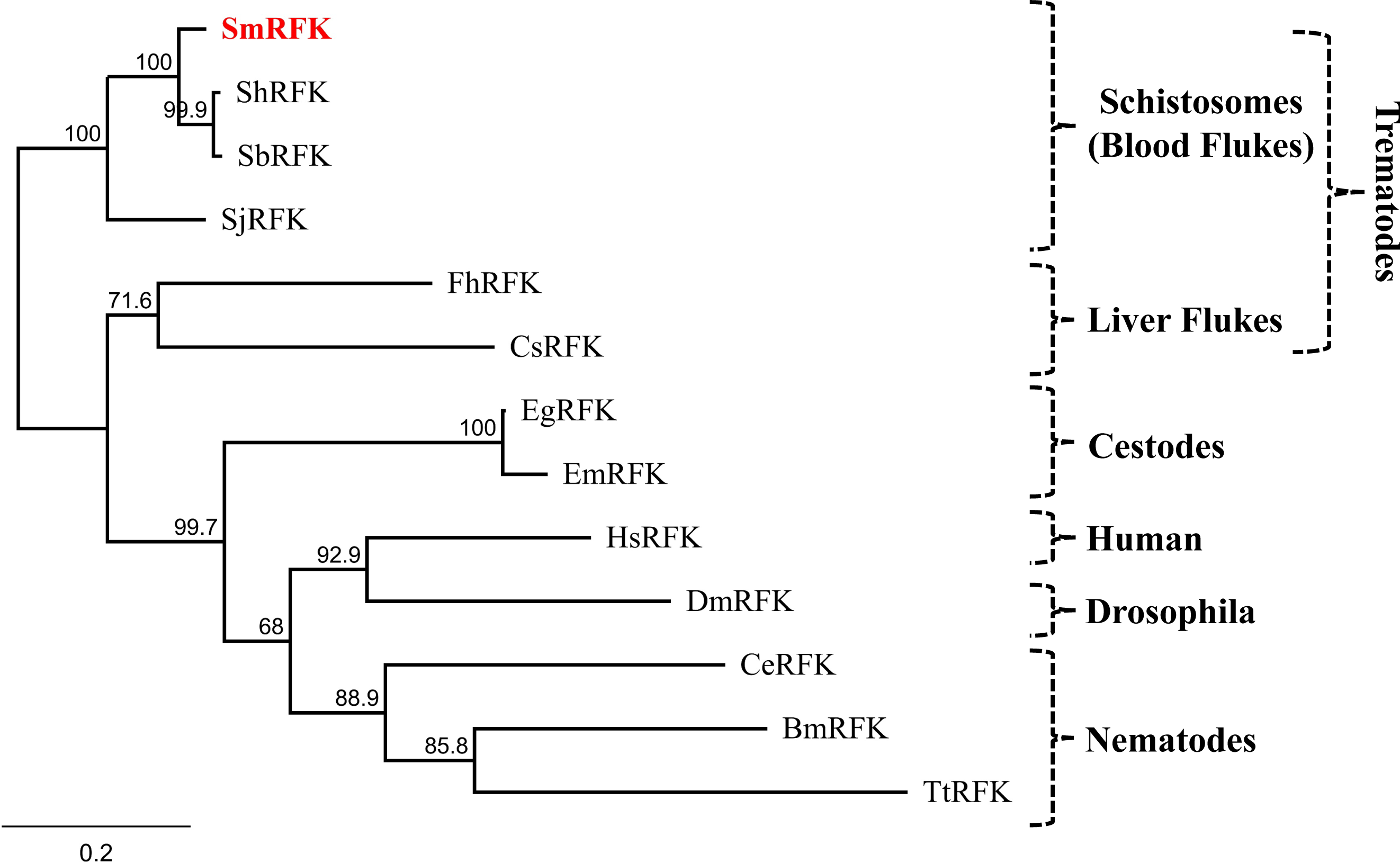
A. Alignment of SmRFK with other members of the riboflavin kinase (RFK) protein family. Multiple sequence alignment was generated using Clustal Omega. Highly conserved residues are indicated with (*), amino acids with strongly similar properties are indicated with (:), amino acids with weakly similar properties are indicated with (.). Residues that are involved in flavin binding are colored red and indicated with ($); residues involved in ADP/ATP binding are colored blue and indicated with (#), and the conserved Thr^32^ residue involved in magnesium ion binding is colored green and indicated with (↓). The conserved ^31^PTAN^34^ motif of RFK protein family members is boxed. **B.** Phylogenetic tree generated by Neighbor-Joining of selected RFKs generated using Geneious Prime Software. Numbers on the branches reflect the consensus branch support value (%), and the scale bar indicates substitutions per site. Abbreviations, species names, and GenBank accession numbers are as follows: Sm, *Schistosoma mansoni* (XP_018647028.1); Sh, *Schistosoma haematobium* (KAH9580372.1); Sb, *Schistosoma bovis* (RTG86096.1); Sj, *Schistosoma japonicum* (AAW27530.1); Fh, *Fasciola hepatica* (THD21035.1); Eg, *Echinococcus granulosus* (XP_024352885.1); Hs, *Homo sapiens* (NP_060809.3); Ce, *Caenorhabditis elegans* (NP_501922.1); Dm, *Drosophila melanogaster* (AAL28446.1); Bm, *Brugia malayi* (XP_042930421.1); Tt, *Trichuris trichiura* (CDW52326.1); Em, *Echinococcus multilocularis* (CDS43726.1); Cs, *Clonorchis sinensis* (KAG5454931.1).

A phylogenetic tree illustrating the relationships between RFK homologs is depicted in figure 5B, where SmRFK is highlighted by red text (top). Clearly, RFK sequences from schistosome species are most closely related and these, along with homologs from other trematodes and cestodes form a platyhelminth RFK clade.

In other systems, an FAD synthase (FADS) enzyme converts FMN to FAD and, by blasting the human FADS protein sequence against schistosome sequence databases, we identified a clear *S. mansoni* FADS homolog that appeared not to be full length by comparison with other available FADS sequences. 5’RACE was employed to successfully amplify the amino terminal coding DNA, and this permitted the assembly of a full-length sequence designated SmFADS (accession number OR495728). This protein is predicted to contain 547 amino acids with a predicted molecular mass of 62,191 Daltons and a pI of 6.15. Figure 6A shows an alignment of the SmFADS protein with counterparts from *S. haematobium* and *S. japonicum* as well as homologs from a variety of other organisms. Similar to human and other eukaryotic FADS, SmFADS also contains two major domains: an N-terminal molybdenum cofactor (MoCF) binding domain (highlighted in yellow in figure 6A), and a C-terminal phosphoadenosine phosphosulphate (PAPS) domain (so called since it is also found in PAPS reductase enzymes and in PAPS sulphotransferases, highlighted in green in figure 6A) ^31–33^. Highly conserved residues in the MoCF domain that are presumed to bind molybdopterin, are shown in blue and indicated with the **#** symbol (Fig. 6A) ^34^. The SmFADS PAPS catalytic domain exhibits five conserved structural motifs that are involved in ATP and Mg^2+^ binding (black boxed residues in Figure 6A)^35^. Those motifs include an ADE (adenine binding) motif, two arginine containing motifs (Arg-1 and Arg-2), a γ-phosphate motif and a pyrophosphate loop (PP-loop) ^35^. A flavin binding motif is boxed in red. Specific conserved residues known to be involved in substrate/product interactions in the PAPS domain are indicated by the $ symbol in figure 6A ^36^.

**Figure 6.**
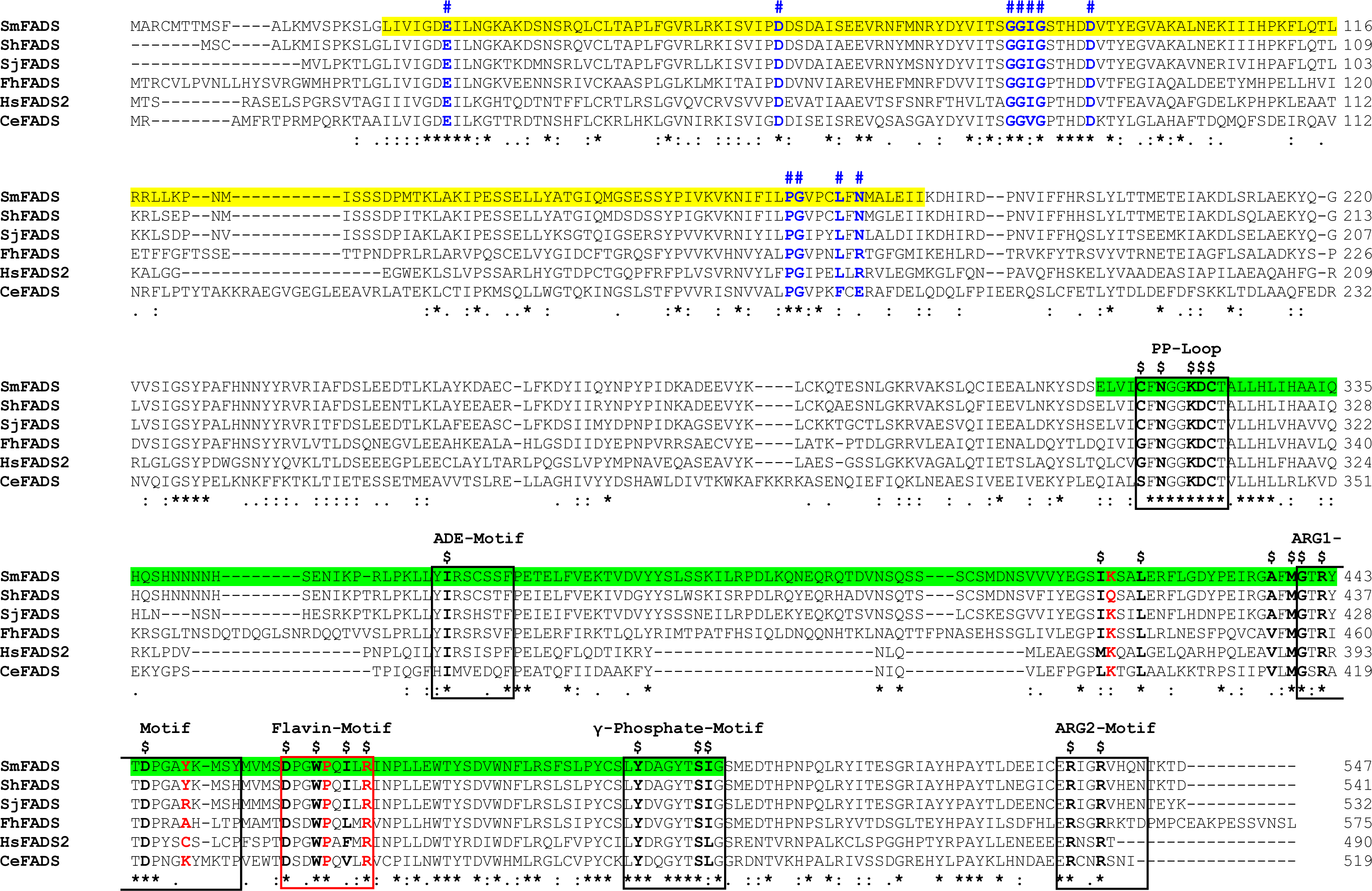
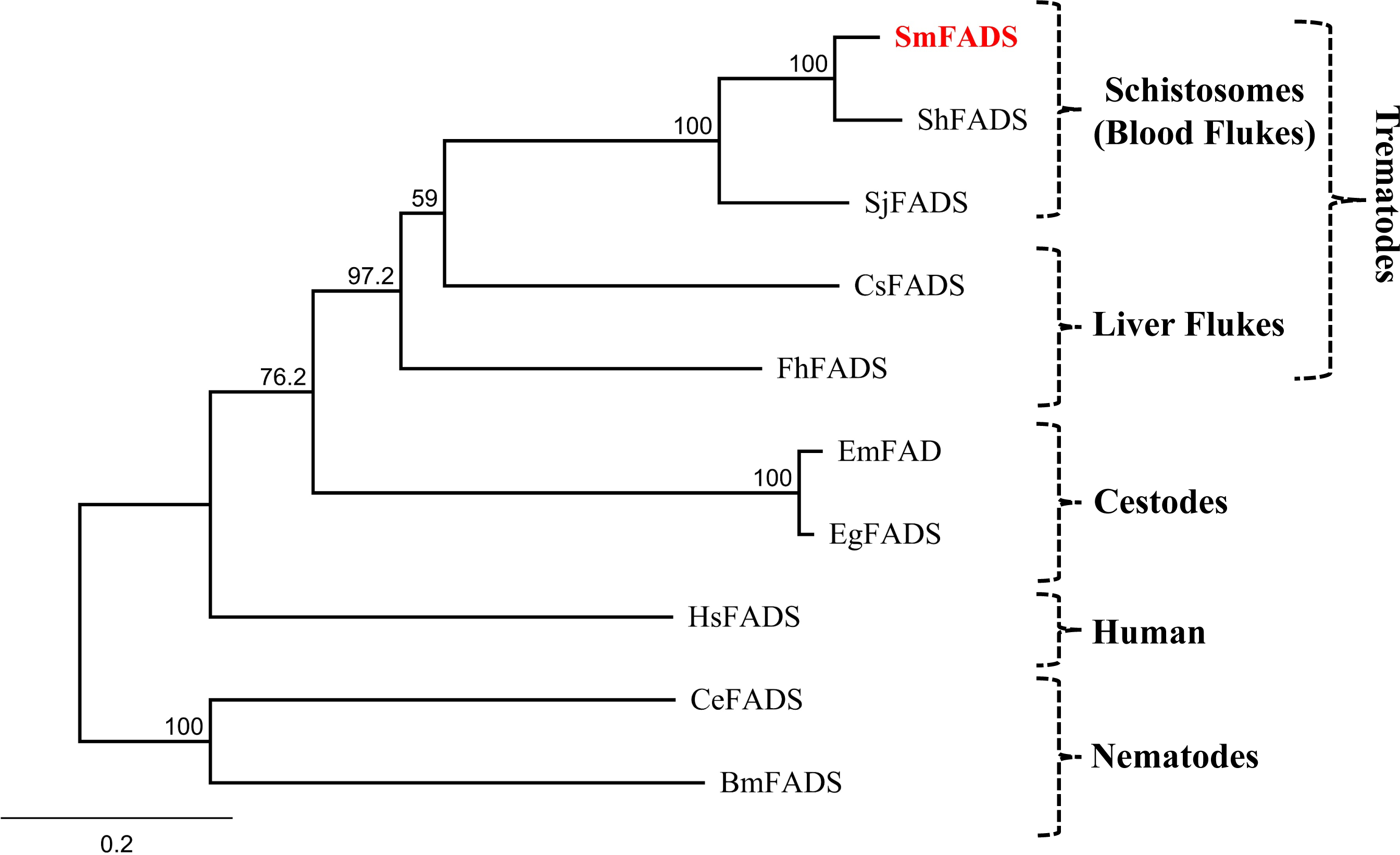
A. Alignment of SmFADS with other members of the FAD synthetase (FADS) protein family. Multiple sequence alignment was generated using Clustal Omega. The SmFADS protein contains 2 major domains: an N-terminal molybdenum cofactor (MoCF) binding domain (MoCF-BD) (L^22^-I^182^; highlighted in yellow in SmFADS) and a C-terminal phosphoadenosine phosphosulphate (PAPS) domain (E^312^–G^501^; highlighted in green in SmFADS). Residues conserved in all six sequences are indicated with (*), amino acids with strongly similar properties are indicated with (:), amino acids with weakly similar properties are indicated with (.). Conserved residues in MoCF domain that are hypothesized to bind molybdopterin are in blue and indicated with (#). Five conserved structural motifs of SmFADS found in the PAPS catalytic domain are bounded by black boxes and named. A flavin binding motif is boxed in red. Residues involved in substrate/product interaction in the PAPS domain are indicated by ($). Key conserved residues in the enzyme’s active site are in red. **B.** Phylogenetic tree generated by Neighbor-Joining of selected FADS enzymes generated using Geneious Prime Software. Numbers on the branches reflect the consensus branch support value (%), and the scale bar indicates substitutions per site. The abbreviations, species names, and GenBank numbers of the organism used in the multiple sequence alignment (A) in the phylogenetic tree (B) analyses are as follows: Sm, *Schistosoma mansoni* (WMM48042.1); Sh, *Schistosoma haematobium* (KAH9594999.1); Sj, *Schistosoma japonicum* (KAH8853370.1); Fh, *Fasciola hepatica* (THD26248.1); Hs, *Homo sapiens* (NP_958800.1); Ce, *Caenorhabditis elegans* (NP_001022286.1); Cs, *Clonorchis sinensis* (GAA53108.1); Bm, *Brugia malayi* (XP_001893012.2); Eg, *Echinococcus granulosus* (CDS20212.1); Em, *Echinococcus multilocularis* (CDI98557.1).

A phylogenetic tree illustrating the relationships between FADS homologs is depicted in figure 6B, where SmFADS is highlighted by red text. As expected, the predicted FADS sequences from all schistosome species are closely related and all platyhelminth sequences cluster together. As was seen in the case of RFK homolog analysis earlier, the nematode FADS species sequences are the most evolutionarily distinct from the schistosome sequences, among all sequences compared here.

## Discussion

When schistosomes are incubated in murine plasma they alter its metabolomic profile; the level of FAD decreases while the level of FMN increases. There is no significant change in the levels of these metabolites in control plasma that does not contain worms. The changes seen in the presence of worms are best understood if the parasites have an ability to cleave any FAD in the plasma leading to the generation of FMN. In human cells it has been reported that exogenous FAD is hydrolyzed by the ecto-5’ nucleotidase CD73 to FMN ^16^. We hypothesized that in schistosomes an equivalent tegumental ecto-enzyme could perform the same function. However, multiple analyses of the proteomic composition of the schistosome tegument, failed to identify a 5’nucleotidase homolog ^17,27,28^. However, we have characterized a GPI-linked schistosome tegumental phosphodiesterase/pyrophosphatase designated SmNPP5 that can cleave a related metabolite – NAD ^20^. SmNPP5 can additionally hydrolyze ATP, ADP and ADPR ^21^. SmNPP5 is an essential enzyme for schistosomes since parasites whose SmNPP5 gene has been suppressed using RNA interference cannot establish a robust infection in experimental animals ^19^. To test the hypothesis that SmNPP5 can also hydrolyze FAD, we incubated purified rSmNPP5 with FAD and successfully detected reaction product (FMN) generation over time. Adding calf intestinal phosphatase (cip) to the reaction products resulted in the release of Pi, an outcome also consistent with the cleavage of FAD to FMN. We also showed directly that live parasites could cleave extracellular FAD and that parasites whose SmNPP5 gene was suppressed by RNAi were significantly impaired in their ability to cleave FAD compared to controls. This proves that SmNPP5, present at the surface of intravascular schistosomes, can indeed cleave exogenous FAD.

FAD is a key coenzyme that are required for catalytic activity of several enzymes including Interleukin 4 Induced 1(IL-4I1), a glycosylated protein that is secreted from myeloid cells as well as B and T cells ^9,10^. IL-4I1 belongs to the L-amino-acid oxidase (LAAO) family of flavoproteins; it performs oxidative deamination of phenylalanine into phenylpyruvate, liberating H_2_O_2_ in the process. Since schistosomes can be efficiently killed by H_2_O_2,_ ^29,30^, we reasoned that SmNPP5-mediated cleavage of coenzyme FAD might block the action of IL-4I1. By preventing the generation of H_2_O_2_ around them, cleaving FAD to inhibit IL-4I1 action might be selectively advantageous for the worms. To test this, we incubated IL-4I1 with rSmNPP5 and compared IL-4I1’s activity with that seen when IL-4I1 was incubated with a control protein (BSA). There was no significant difference seen in the activity of IL-4I1 in either case, showing that SmNPP5 could not block IL-4I1 action. We conclude that in this experiment, FAD, covalently bound to IL-4I1 ^9^, is inaccessible to SmNPP5. Perhaps selection favors covalently binding essential factors like FAD to some enzymes specifically to prevent coenzyme destruction by pathogen produced FADase enzymes like SmNPP5.

In work with human cells, exogenous FMN can be hydrolyzed by an ecto-alkaline phosphatase to produce riboflavin ^16^. Schistosomes express a well-characterized ecto-alkaline phosphatase (SmAP) that has been shown to dephosphorylate several substrates including AMP, CMP, GMP and TMP as well as the bioactive lipid sphingosine-1-phosphate (S1P) and the proinflammatory/prothrombotic polymer, polyphosphate (polyP) ^25,26^. SmAP has also been shown to contribute to the metabolism of another vitamin, pyridoxal phosphate (PLP, vitamin B6) ^37^. We have shown that SmAP can dephosphorylate PLP to generate pyridoxal and we speculate that this permits the efficient uptake of this vitamin by the worms ^37^. Likewise, we hypothesize here that SmAP-mediated dephosphorylation of FMN to yield riboflavin helps to generate a pool of this vital metabolite around the worms. Working with human cells, it has been shown that cleavage of FMN by an ecto-alkaline phosphatase generates riboflavin ^16^ and we hypothesized that the same scenario applies in the case of intravascular schistosomes. In work described here, we have shown that rSmAP can indeed cleave FMN to generate riboflavin (and free phosphate). In addition, we showed that live parasites can cleave extracellular FMN and that parasites whose SmAP gene has been suppressed by RNAi are significantly impaired in their ability to cleave FMN compared to controls. This proves that SmAP, present at the surface of intravascular schistosomes, can indeed cleave exogenous FMN to generate vitamin B2. If SmAP on worms cleaves extracellular FMN to generate riboflavin, we would predict that riboflavin levels would increase with time in plasma containing worms, but we do not see this. Instead, while FAD and FMN levels change in plasma containing schistosomes, plasma riboflavin levels remain essentially unchanged. Perhaps this reflects the uptake of any additional riboflavin generated by SmAP-mediated FMN cleavage by the worms, leaving basal plasma levels relatively unchanged? Note that the AMP (generated along with FMN) by SmNPP5-mediated cleavage of FAD can also be dephosphorylated by SmAP to generate adenosine (and phosphate) ^25^. This is noteworthy because schistosomes cannot synthesize purines *de novo* ^38^. The worms must take in chemicals such as adenosine from their environment to survive, in addition to vitamins such as riboflavin.

Riboflavin generated in the vicinity of intravascular worms via the pathway just mentioned could be taken into the parasites via the recently described *S. mansoni* riboflavin transporter protein SmaRT ^15^. SmaRT is a 531 amino acid protein that has been shown to be capable of mediating riboflavin uptake following its heterologous expression in CHO-S cells ^15^. Uptake is sodium independent and occurs over a wide pH range. The protein is expressed in the tegument and widely in the internal tissues of adult schistosomes ^15^. Thus, SmaRT is likely important both for directly importing riboflavin from host blood and for distributing the vitamin throughout the body of the parasites.

Much of the riboflavin taken up by schistosomes would need to be converted back into both FMN and FAD, in order to generate sufficient levels of these vital coenzymes for efficient parasite metabolism. To explore the possibility that schistosomes possess enzymes that could drive FMN and FAD synthesis from imported riboflavin, we queried schistosome sequence databases for the presence of homologs of human enzymes that promote these reactions. In this manner, a 154 amino acid schistosome protein – SmRFK belonging to the RFK protein family, was identified that possesses strongly conserved motifs known to be involved in flavin and ATP binding. Likely, SmRFK could convert riboflavin to FMN inside schistosomes.

Analysis of schistosome sequence databases further identified a 547 amino-acid schistosome protein with strong homology to members of the FAD synthetase family which we designate SmFADS. Like other eukaryotic FAD synthetases, SmFADS has two major domains: an N-terminal molybdenum cofactor (MoCF) binding domain, and a C-terminal phosphoadenosine phosphosulphate (PAPS) domain. The SmFADS PAPS domain contains all motifs and residues known to be required for FAD synthesis and therefore most likely acts to convert FMN to FAD inside schistosomes. Figure 7 summarize our major findings: starting at the top right, external FAD can be cleaved to FMN (and AMP) via the GPI-linked ectoenzyme SmNPP5 (pink box). FMN can be cleaved to riboflavin (RF and phosphate) by the GPI-linked ectoenzyme SmAP (yellow box). Riboflavin can then be imported into schistosomes via the riboflavin transporter, SmaRT (yellow cylinder, top left). Once internalized, the riboflavin could be converted back to FMN via SmRFK (bright blue box) and some FMN could be converted to FAD via SmFADS (green box). Both of these reactions require ATP.

**Figure 7.**
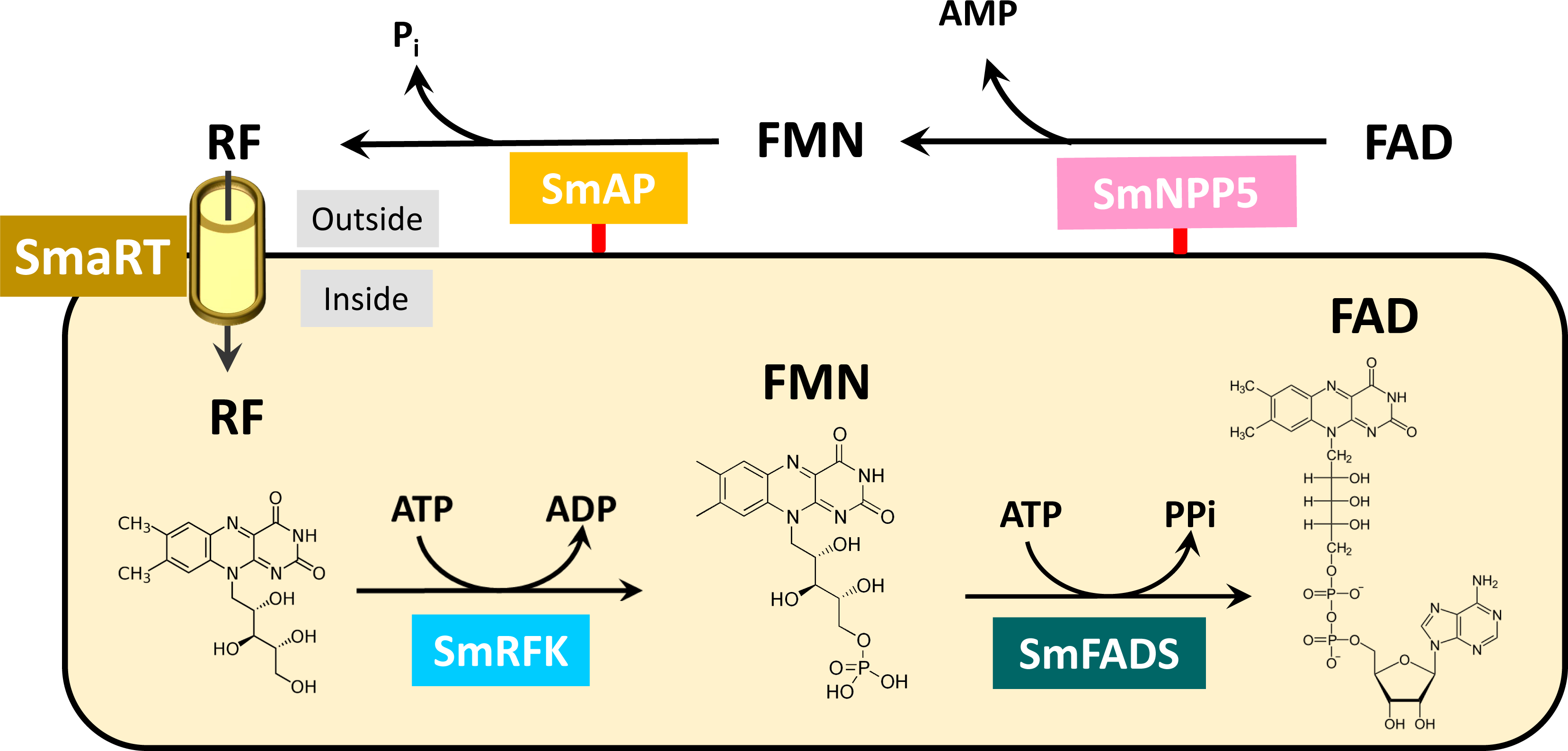
Vitamin B2 metabolism in schistosomes. Ectoenzyme SmNPP5 (depicted at top right, pink box) can cleave FAD to generate FMN (and AMP). Ectoenzyme SmAP (bright yellow box) can cleave FMN to generate RF (and Pi). RF can be imported into the worms via the transporter protein SmART (yellow cylinder, top left). Depicted in the yellow rectangle is the hypothetical biochemical pathway within *S. mansoni* leading to the regeneration of FMN by the action of SmRFK (bright blue box) and then to the regeneration of FAD by the action of SmFADS (dark green box). The chemical structures of RF, FMN and FAD are shown. The red lines depicted at the base of SmNPP5 and SmAP represent GPI-linkages to the external schistosome plasma membrane.

It has been reported that riboflavin can inhibit platelet aggregation *in vitro* ^39^. Thus, it is possible that the actions of SmNPP5 and SmAP to increase the extracellular pool of vitamin B2 around intravascular worms could have a benefit beyond simply supplying the parasites with an easy source of the vitamin. If excess riboflavin can also help to control thrombus formation near the parasites, generating higher levels of extracellular riboflavin by the worms could be even more selectively advantageous.

This work extends the known substrates for the schistosome ectoenzyme SmNPP5 to now include FAD. Similarly, the list of known substrates for SmAP extends to FMN. These two schistosome tegumental ectoenzymes can clearly exert a profound impact on the biochemistry of the worm’s local environment. Given their expanding list of known substrates, it is clear that these enzymes can have a great impact especially on the set of purine-containing metabolites around the worm – collectively known as the worm’s purinergic halo ^40^. We have argued that in some instances the enzymes act to destroy host purinergic signaling molecules that could otherwise drive worm-damaging immune or hemostatic reactions ^40,41^. Here we argue that the enzymes also function to generate the vital metabolite riboflavin around the worms. Blocking SmAP and/or SmNPP5 using chemotherapy or by immunological means should help deprive the worms of essential nutrients and could help to debilitate them and so control infection.

## Methods

### Parasites and Mice

The Puerto Rican strain of *Schistosoma mansoni* was used. Adult male and female parasites were recovered by perfusion from Swiss Webster mice that were infected with ∼100 cercariae, 7 weeks previously ^42^. Schistosomula were prepared from cercariae released from infected snails. All parasites were cultured in DMEM/F12 medium supplemented with 10% heat-inactivated fetal bovine serum, 200 µg/ml streptomycin, 200 U/ml penicillin, 1 µM serotonin, 0.2 µM Triiodo-l-thyronine, 8 µg/ml human insulin and were maintained at 37°C, in an atmosphere of 5% CO_2_ ^25^. Protocols involving animals were approved by the Institutional Animal Care and Use Committees (IACUC) of Tufts University.

### FAD, FMN and Riboflavin Detection in Murine Plasma

Blood was collected from the tail veins of 10 Swiss Webster mice into heparinized tubes. Blood cells were pelleted by brief centrifugation and the plasma generated was pooled and aliquoted. Adult schistosomes (∼50 pairs) were incubated in one 500 µl murine plasma aliquot which was incubated at 37°C. A control aliquot (without worms) was similarly treated. Samples, collected at baseline (0 min) and after 20- and 60-min incubation with or without parasites, were subjected to metabolomic analysis at Metabolon Inc. The relative levels of FAD, FMN and riboflavin are described; these are extracted from a global metabolomics analysis carried out using the pipeline developed by Metabolon. At least 4 samples per treatment/time point were tested. Briefly, each plasma sample was prepared by solvent extraction and the resulting extract was split into equal parts and then applied to gas chromatography/mass spectrometry (GC/MS) and liquid chromatography tandem MS (LC/MS/MS) platforms ^43^. FAD, FMN and riboflavin were each identified by their retention time and mass by comparison to purified standards. Results are expressed relative to the baseline measurement (0 min), set at 1.

### FAD Hydrolysis by rSmNPP5

Recombinant SmNPP5, expressed in CHO-S cells, was prepared as described [20, 24, 30]. To assess its ability to cleave FAD, 0.1µg rSmNPP5 was added to a solution of FAD (50µM or 500µM) in assay buffer (50mM Tris-HCl, pH 9.0, 120mM NaCl, 5mM KCl, 50mM glucose, 2mM CaCl_2_). Fluorescence associated with generated product (FMN) was measured continuously over time at 375 nm excitation and 520 nm emission using a Synergy HT microplate reader (Bio-Tek Instruments). In parallel experiments, aliquots of the reaction mixture were also recovered at 0.5, 1, 2, and 5 h; calf intestinal phosphatase (cip, 70U/ml) was added to each aliquot for 30 min and any inorganic phosphate that was generated was measured using a commercial Phosphate Colorimetric Assay Kit (BioVision), following the manufacturer’s instructions. The Michaelis–Menten constant (Km) of rSmNPP5 for FAD was determined using standard assay conditions described above, with varying concentrations of FAD (0-800 µM). Standard curves were generated using known amounts of FMN. The Km for FAD was determined from these data using GraphPad Prism V9.0 (GraphPad Software).

### IL-4I1 Assay

The effect of rSmNPP5 on IL-4I1 activity was assessed using a commercial coupled horseradish peroxidase (HRP) assay, according to the manufacturer’s instructions (R&D Systems). Briefly, 0.1µg rIL-4I1, 1mM phenylalanine (as substrate), 1 unit/ml HRP and 50µM Amplex Ultra Red with either 1µg rSmNPP5 or BSA (as control) were added in 100µl final volume of assay buffer (50 mM sodium phosphate, pH 7.0). Each experiment included a substrate only and rSmNPP5 only control (i.e., lacking IL-4I1, as additional controls). The assays were performed in black, clear bottomed 96 well plates and fluorescence was read at excitation and emission wavelengths of 544 nm and 590 nm, respectively in kinetic mode, every 5 minutes, for 1 h using a Synergy HT microplate reader (Bio-Tek Instruments).

### FMN Hydrolysis by rSmAP

To look for the ability of rSmAP to dephosphorylate FMN, 1µg enzyme was added to 500 µl assay buffer (50 mM Tris–HCl, pH 9.0, 5 mM KCl, 135 mM NaCl, 10 mM glucose, 10 mM MgCl_2_) containing 1 mM FMN. Any phosphate generated following FMN cleavage was measured at selected time points using a Phosphate Colorimetric Assay Kit, as above.

Thin layer chromatography (TLC) was employed as an additional method of monitoring SmAP-mediated FMN cleavage. To visualize enzyme reaction products, 1µg rSmAP was first incubated with 1 mM NMN in assay buffer, as above. After 1h, aliquots from these reactions (and the equivalent volumes of chemical standards (FMN and RF) diluted to 1 mM in assay buffer), were spotted ∼1 cm from the base of a TLC LC Silica gel 60 F245 aluminum sheet (20L×L20 cm, EMD Millipore). Spots were allowed to dry, and separation of their chemical constituents was achieved using a mobile phase composed of n-butanolL:Lacetone : acetic acid (glacial)L:Lammonia (5%) : water (45L:L15L:L10L:L10L:L20). Analytes were visualized under UV at 254 nm [36].

### FAD and FMN Hydrolysis by Living Schistosomes

To monitor FAD and FMN hydrolysis by living schistosomes, schistosomula (in groups of ∼2,000) were incubated in 200 µl assay buffer (50 mM Tris-HCl buffer (pH 9), 5 mM KCl, 135 mM NaCl, 5 mM KCl, 10 mM glucose, 2mM CaCl_2_, 10 mM MgCl_2_) containing either 2 mM FAD or FMN. To monitor FAD cleavage, changes in fluorescence were monitored over time, as described above. To monitor FMN cleavage, Pi levels were monitored at different timepoints, as above.

### Ectoenzyme Gene Suppression using RNAi

Adult worms were electroporated with either an siRNA (10 µg) targeting SmNPP5 (SmNPP5 siRNA 1: 5L-TTGATGGATTTCGTTATGATTACTTTG-3L) or SmAP (SmAPsiRNA1: 5’-AAGAAATCAGCAGATGAGAGATTTAAT-3’), or with a control siRNA that targeted no sequence in the schistosome genome (Control: 5’-CTTCCTCTCTTTCTCTCCCTTGTGA-3’) or with no siRNA, as described previously ^19,23^. To assess the level of target gene suppression, RNA was isolated from some worms from each group two days later using the TRIzol Reagent (Thermo Fisher Scientific, MA), as per the manufacturer’s guidelines. Residual DNA was removed by DNase I digestion using a TurboDNA-free kit (Thermo Fisher Scientific, MA). cDNA was synthesized using 1 µg RNA, an oligo (dT)_20_ primer and Superscript III RT (Invitrogen, CA). Gene expression of SmNPP5 or SmAP was measured by quantitative real time PCR (RT-qPCR), with custom TaqMan gene expression systems from Applied Biosystems, CA using primers and probes, as previously ^19,23^. Alpha tubulin was used as the endogenous control gene for relative quantification, as described ^44^, employing the ΔΔCt method ^45^. Results obtained from parasites treated with the irrelevant, control siRNA were used for calibration. For graphical representation, 2^-ΔΔCt^ values were normalized to controls and expressed as percent difference. Seven days post siRNA treatment, the ability of gene suppressed v control parasite groups (in replicate) to cleave FAD or FMN were compared using the assays described above.

### Cloning cDNAs Encoding Cytosolic Vitamin B2 Metabolizing Enzymes of Schistosomes

To look for homologs in *S. mansoni* of the human enzymes that are central to vitamin B2 metabolism within cells, we used the human riboflavin kinase sequence (GenBank accession number Q969G6) and the human NAD synthase sequence (Q5T196) to blast against all available “schistosomatidae” family sequences at NCBI

(https://blast.ncbi.nlm.nih.gov/Blast.cgi?PAGE=Proteins). Clear sequence homologs were identified in each case and were designated SmRFK and SmFADS, respectively.

Primers (SmRFK**-**F: 5’-ATGTTTGTTAATTTAACAGCTGGAGC-3’and SmRFK-R: 5’-TCATCGGTCTTTTTCATTGAATAAGTCATG-3’) were generated and used in a PCR, with adult worm cDNA as template, to amplify the complete SmRFK coding sequence. The PCR product was purified, cloned into pCRII-TOPO and transformed into TOP10 *E. coli* (Thermo Fisher Scientific, MA), using standard techniques. Plasmid was purified from a selection of recombinant transformants, and plasmid DNA inserts were sequenced at Genewiz, Inc. MA.

The SmFADS sequence appeared to be incomplete in comparison with its human homolog. Therefore, a SMARTer^®^ RACE 5’/3’ Kit was employed to identify the complete 5’-end of SmFADS, following the manufacturer’s instructions (Takara Bio). This approach resulted in the identification of both SmFADS ends. To confirm these results, the entire open reading frame of SmFADS was amplified as one unit by PCR using the primers FADS-F: 5’-ATGGCGCGGTGTATGACTACTATGTC-3’, and FADS-R:5’-TCAATCAGTTTTAGTATTTTGATGC-3’.

Multiple protein sequence alignment was performed using the online Clustal Omega tool (https://www.ebi.ac.uk/Tools/msa/clustalo/). Phylogenetic trees were constructed by Neighbor-Joining with Jukes-Cantor genetic distance mode, using Geneious Prime Software (Biomatters Ltd.). Trees were reassembled using the Bootstrap method, with 1000 replicates.

### Statistical Analysis

Data are presented as Mean ± SD. Means were compared by t-test (two-tailed, unpaired) for comparison of two groups or by one-way ANOVA for comparison of more than two groups using GraphPad Prism 10.0 (GraphPad Software). For metabolite comparisons, Welch’s two-sample t-test was used to identify biochemicals biomolecules that differed significantly between groups. P<0.05 was considered significant.

## Acknowledgements

This work was funded with support from NIH-NIAID grant AI056273. Infected snails were provided by BRI via the NIAID schistosomiasis resource center under NIH-NIAID Contract No. HHSN272201700014I.

